# Polar organization of H&E dyes in histology tissue revealed by polarimetric nonlinear microscopy

**DOI:** 10.1101/2025.05.19.654795

**Authors:** Mykolas Maciulis, Viktoras Mazeika, Lukas Kontenis, Danielle Tokarz, Richard Cisek, Danute Bulotiene, Vitalijus Karabanovas, Virginijus Barzda

## Abstract

Structural organization of harmonophores used in hematoxylin (H) and eosin (E) staining is studied with polarimetric multimodal second-harmonic generation (SHG), third-harmonic generation (THG) and multiphoton excitation fluorescence (MPF) microscopy in rat tail tendon histology sections. The polarimetric microscopy imaging reveals that hemalums (complexes of hematoxylin and aluminum) are well aligned with *C*_6*h*_ symmetry along the collagen fibers in H-stained tissue, while eosin Y is partially aligned along the fibers in E-stained tissue and also follows organization of *C*_6*h*_ symmetry. When both hemalum and eosin are used for H&E staining, the dye molecules interact and align noncentrosymmetrically with *C*_6_ symmetry along the collagen fibers, while the stained nuclei appear isotropically organized. The polar alignment of the hemalum and eosin complexes increases the achiral second order susceptibility tensor component ratio 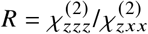 in H&E-stained tissue. The alignment of hemalum and eosin molecules, and their complexes in collagenous tissue, must be considered in nonlinear microscopy and polarimetric analysis of H&E-stained histopathology.

## 1. Introduction

Hematoxylin- and eosin-(H&E) stained tissue histopathology is considered a gold standard in cancer diagnostics. Tissue staining with hemalums reveals the nuclear structure, while eosin counter staining highlights proteins in the cells and the extracellular matrix (ECM) [1, 2]. ECM counterstained with eosin has low specificity in white light microscopy, and is usually not utilized as a marker for cancer diagnostics. In contrast, ECM collagen structure can be examined by second-harmonic generation (SHG) microscopy with high specificity [3–5] and represents a sensitive biomarker for diagnostics of cancer [6–11]. Furthermore, polarization-resolved SHG imaging can be utilized to investigate the ultrastructure of collagen within the focal volume of a microscope [12, 13]. Therefore, polarimetric SHG microscopy can serve as a sensitive tool to characterize structural changes in ECM with applications in cancer diagnostics [10, 14–16].

Nonlinear optical microscopy has emerged as a powerful method for imaging of biological tissues [17–19]. Multimodal nonlinear microscopy can provide simultaneous imaging with multiphoton excitation fluorescence (MPF), SHG, and third-harmonic generation (THG) contrast mechanisms [20]. MPF originates mostly from eosin in H&E-stained tissues [21] with some contribution from tissue autofluorescence [22, 23]. Eosin stains proteins non-specifically and highlights tissue extracellular matrix (ECM) as well as cytoplasm [1, 2]. In contrast, SHG does not require staining and originates from noncentrosymmetric structures, predominantly from collagen [4] and myosin [24]. Collagen is a fibrillar protein comprised of a hierarchical noncentrosymmetric structure consisting of tripple helixes that self assemble into fibrills, and, in turn, into fibers. Collagen fibers have polar structure that can be revealed by polarimetric SHG (P-SHG) microscopy [25]. H&E staining is reported to affect SHG intensity and the second order susceptibility tensor elements of collagen [26–28]. Therefore, it is important to decipher the molecular mechanisms for the modification of SHG in H&E-stained tissues. THG occurs at structural interfaces with local changes of refractive index or third-order nonlinear susceptibility [29, 30]. Focused beams do not produce THG in homogeneous media due to Gouy phase shift [31], but generate a signal from nanostructures and interfaces. THG has been shown to originate from hematoxylin aggregates accumulated in the nuclei [21]. However, lower intensity signals originating from the tissue stroma remained unaddressed in THG images.

The interaction of the H&E stains with the biological tissue is complex [26, 28]. In this study, we apply polarimetric nonlinear microscopy to investigate the organization of hemalum and eosin Y molecules bound to the tissue during the staining procedure. The alignment of stain molecules has a profound effect on the MPF, SHG and THG signals. These effects are studied in H-, E- and H&E-stained histology sections of rat tail tendon (RTT) tissue. The results show that, when stained with one of the dyes, hemalums and eosin distribute with *C*_6*h*_ cylindrical symmetry having the symmetry axis along the collagen fibers. Therefore, SHG is not modified by the staining. In contrast, when both stains are used, they interact and noncentrosymmetrically distribute with *C*_6_ cylindrical symmetry axis along the collagen fibers in the H&E-stained tissue. The alignment of hemalum and eosin molecules in the ECM modifies the SHG response by decreasing the signal intensity and increasing the achiral susceptibility ratio 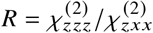 of the H&E-stained RTT tissue. Therefore, the effects of H&E staining on the MPF, SHG and THG imaging have to be taken into account for the histopathology investigations.

## 2. Materials and methods

### 2.1. Nonlinear microscope setup

Nonlinear microscopy is performed using a home-built polarimetric laser-scanning microscope which can collect MPF, SHG and THG signals. A femtosecond oscillator (FLINT FL1, Light Conversion) generating 100 fs pulses at a 76 MHz pulse repetition rate and a central wavelength of 1030 nm is used for excitation. Images are obtained by raster-scanning the laser beam using a set of galvo mirrors (Saturn 5B, Pangolin). Incident and outgoing polarization states are controlled and analyzed with a polarization state generator (PSG) and analyzer (PSA), respectively. The PSG is comprised of a fixed polarizer (LPNIR100, Thorlabs), followed by motorized zero-order half- and quarter-wave plates (WPH10M-1030 and WPQ10M-1030, Thorlabs). The PSA contains a broadband wire grid polarizer (WP25M-UB, Thorlabs) for THG, or motorized zero-order half- and quarter-wave plates (WPH10M-514 and WPQ10M-514, Thorlabs), followed by a fixed polarizer (LPVISA100, Thorlabs) for SHG. A 20× 0.75 numerical aperture (NA) objective (20×/0.75 Plan Apo Lambda, Nikon) is used for excitation. The nonlinear signals are collected in the transmission direction using a 0.45 NA singlet lens and detected with a photomultiplier tube (PMT) (H10682-210, Hamamatsu) operating in photon-counting mode. MPF, SHG and THG signals are separated by placing filters in front of the PMT: a 550 nm longpass filter (FELH0550, Thorlabs) and a 750 nm shortpass filter (FESH0750, Thorlabs) for MPF, a 10 nm band-pass filter (BP515-10, Edmund Optics) at 515 nm for SHG, and a 10 nm band-pass filter (FBH343-10, Thorlabs) at 343 nm for THG. Scan control and image acquisition are accomplished using a data acquisition card (PCIe-6353, National Instruments). Microscope control and image acquisition are performed with a custom program written in LabVIEW. The polarization independent intensity images (Fig. 1) are constructed from the sum of four images obtained with different incident linear polarization states oriented at -45, 0, 45 and 90 degrees with respect to the vertical axis of the image, and no analyzer. The images are obtained with a 7.8 µs pixel dwell time. The average imaging power was 10 mW at the sample.

**Fig. 1.**
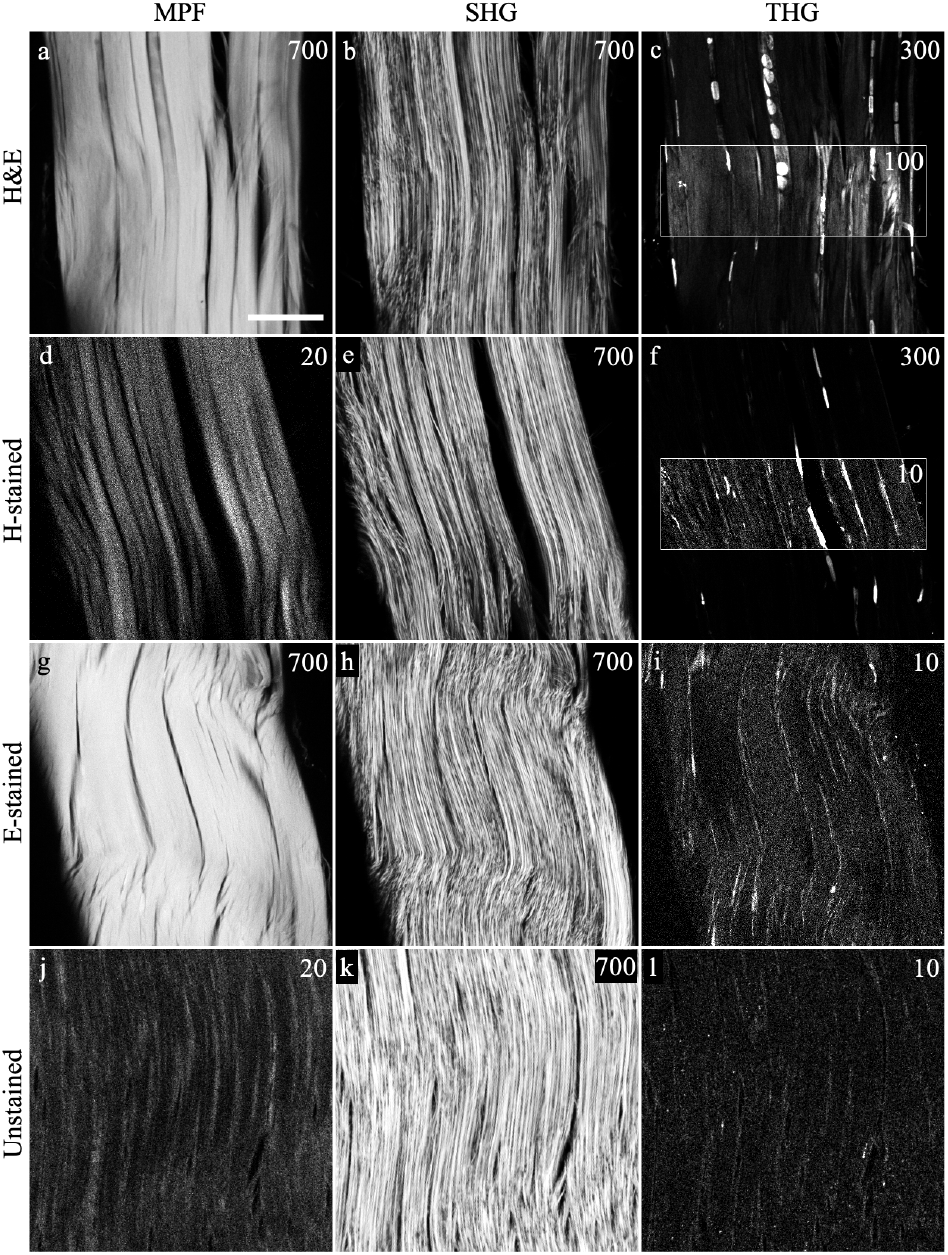
Multimodal nonlinear images of RTT tissue stained with H&E (a-c), hematoxylin (d-f), eosin (g-i), and unstained (j-l). MPF (a, d, g, j), SHG (b, e, h, k) and THG (c, f, i, l) images are shown in corresponding columns. Inset boxes have the upper count limit lowered to show the lower THG signal distribution, while high intensity values from the nuclei are saturated (c, f). Scale bar 50 µm. The images contain 400 × 400 pixels. Maximum photon counts per pixel are given in the upper right corner of each panel. The maximum count value is set so that less than 3% of pixels can exceed it.

### 2.2. Polarimetric measurements of MPF, SHG and THG

The polarization response of the THG (P-THG) is investigated using the polarization-in, polarization-out (PIPO) technique [32], which provides information about the structural organization of harmonophores in biological tissues [33]. The THG polarization-resolved images are collected at 81 combinations of incident and outgoing polarization states for PIPO analysis. The incident and outgoing linear polarization states are rotated over 9 different orientations evenly spaced between 0 and 180 degrees. For each incident polarization state 9 different outgoing polarization states are recorded. Reference frames at the same polarization states are recorded every 9th measurement to check for the sample photostability.

The PIPO analysis is performed on the THG datasets assuming a *C*_6*v*_ symmetry model, with molecular cylindrical axis along *z* direction, and *x, y* perpendicular to the cylindrical axis. Fitting the THG PIPO model to the dataset provides two susceptibility component ratios: 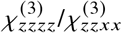 and 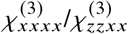. The orientation angle *δ* of the in-image-plane cylindrical axis of the molecular distribution is also obtained from the fit and is defined from the Z-axis of the laboratory coordinate frame, while the Y axis is along the light propagation direction. THG PIPO fitting is performed pixel by pixel with custom program [34]. Polarimetric response of PSA and PSG is calibrated with an isotropic borosilicate glass coverslip for THG and a z-cut quartz wafer for SHG.

The molecular susceptibility SHG tensor achiral component ratio 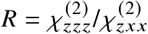 is obtained with double Stokes polarimetry (DSP) as described in [35, 36]. P-SHG measurements are performed with incoming right- and left-circularly polarized (RCP and LCP) light. The polarization of outgoing SHG is probed by collecting linearly polarized (horizontally, vertically and diagonal at ±45 degrees with respect to the laboratory frame Z-axis) as well as RCP and LCP light. The obtained images are used to calculate the Stokes components [36], which in turn are used to calculate the *R* ratio:

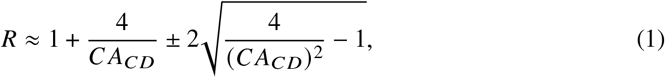

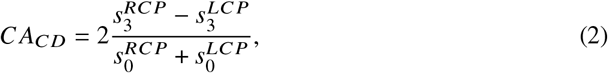

where *C A*_*CD*_ is a circular anisotropy of circular dichroism [37]. The 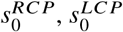 and 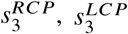 are the SHG Stokes vector components obtained with incoming RCP and LCP light as indicated in the superscript. Kleinman symmetry is assumed in the analysis with 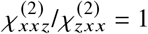.

Linear anisotropy of two-photon absorption (TPA) is determined using fluorescence-detected multiphoton linear dichroism (FDMLD), which is a nonlinear analog of the (linear-excitation) fluorescence-detected linear dichroism (FDLD) [38]. Two-photon excitation fluorescence is detected from 550 nm to 750 nm wavelengths using vertical (V) and horizontal (H) incident linear polarization and FDMLD values are calculated using the pixel intensity *I*_*V*_ and *I*_*H*_:

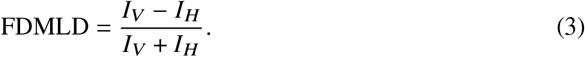

A pixel masking procedure is applied for FDMLD maps, using a threshold of 3 counts for H&E- and E-stained samples, and 1 count for H-stained and unstained samples.

### 2.3. Measurements of the third order molecular hyperpolarizability

To obtain the third order molecular hyperpolarizability value *γ* of eosin Y, *χ*^(3)^ values of 5 solutions of eosin Y disodium salt (E6003, Sigma Aldrich) at different concentrations in distilled water are determined using a home-built nonlinear optical microscopy system described previously [39]. Briefly, the laser source consists of a 1030 nm femtosecond duration pulsed laser (FemtoLux 3, Ekspla), and focusing is achieved using a 10× 0.25 NA air immersion microscope objective lens (Achrostigmat, Carl Zeiss AG). A custom built 0.85 NA collection objective lens (Omex Technologies) is used to collect THG signal in the transmission direction. The THG signal is filtered with a 10 nm band-pass filter (65-129, Edmund Optics), and detected with a photomultiplier tube in photon counting mode (H10682-210, Hamamatsu). By employing an automated translation stage (ASI), capillary tubes (5012, VitroCom) containing the diluted eosin Y solutions are individually translated axially along the laser focal spot. The THG signals generated from the solution-glass and glass-air interfaces closest to the excitation objective lens are recorded at least 10 times per solution.

To calculate the *χ*^(3)^ value of a solution, both the THG intensities at the two interfaces are needed along with the refractive indices of the solution at the laser and third harmonic wavelengths. Therefore, refractive indices of the solutions are measured using a home-built refractometer described previously [39]. Briefly, the refractometer consists of two LED light sources (M340L4 and M1050L2, Thorlabs) centered at wavelengths of 340 nm and 1050 nm, respectively. An optical filter (FLH1030-10, Thorlabs) is used with the 1050 nm LED light to obtain 1030 nm light. The LED light is passed through two yttrium aluminum garnet (YAG) prisms (Red Optronics) which are squeezed together to hold a thin spacer (Seal & Design Canada) and the solution. Transmitted light is detected by a standard CMOS camera (HD Webcam C270, Logitech) while the prisms, spacer and solution are rotated with a motorized rotation stage (CR1-Z7, Thorlabs) until the light diminishes due to the total internal reflection at the YAG-solution interface. The critical angle found for distilled water is used as a reference angle and combined with the critical angles of the eosin Y sample solutions to calculate the refractive indices of the eosin Y solutions at the two wavelengths.

The *χ*^(3)^ values of the eosin Y solutions are calculated as described previously [40] using a *χ*^(3)^ expression by Shcheslavskiy et al. [41] and assuming previously published values for the refractive indices and the *χ*^(3)^ value of Duran borosilicate glass [42]. The *γ* value of eosin Y is extracted from a plot of the *χ*^(3)^ values of eosin Y solutions at different concentrations [40].

### 2.4. Preparation of RTT samples

Albino Wistar female (10-11 weeks old) rats are acquired from the State Scientific Research Institute of the Innovative Medical Center (Vilnius, Lithuania). All animal procedures are carried out in accordance with national and European regulations and are approved by the Lithuanian Animal Care and Use Committee of the State Food and Veterinary Service (Vilnius, Lithuania, approval number G2-156). The excised RTT samples are embedded in paraffin blocks, sectioned longitudinally and deparaffinized. Sections of 5 µm thickness are stained using a standard hematoxylin and eosin staining procedure with Mayer’s hematoxylin and Eosin Y (Leica ST5020 stainer) [43]. When H-stained or E-stained tissue is prepared, the eosin or hematoxylin staining step is omitted, respectively. Mordant is used in all staining procedures except for the unstained sample. Tissue sections are mounted on microscopy slides and covered with coverslips. Four sample types are prepared: H&E-stained, H-stained, E-stained and unstained.

## 3. Results

### 3.1. Multimodal nonlinear microscopy of H and E stained tissue

The effects of H and E staining on the nonlinear optical response in histology tissues are investigated with multimodal MPF, SHG and THG microscopy (Fig. 1). Unstained tendon tissue exhibits relatively low multiphoton excitation autofluorescence (Fig. 1j), while strong SHG is observed, which highlights the structure of collagen fibers (Fig. 1k). The THG signal is very weak in unstained tissue (Fig. 1l) with only a few counts originating from the extracellular matrix. The E-stained tendon tissue shows strong MPF (more than 30 times higher compared to the autofluorescence) due to the high fluorescence yield of eosin (Fig. 1g). The SHG signal from E-stained tissue has similar intensity to the unstained tissue and reveals similar structural features to the unstained tendon (Fig. 1h), while THG intensity is slightly higher than in the unstained tissue (Fig. 1i). THG shows collagen fiber edges and reveals intrafasicular cells. The emergence of THG signal with eosin staining shows that it can be considered as a harmonophore [21]. The third order molecular hyperpolarizability (*γ*) value of eosin Y is found to be 5.5 ± 0.7 × 10^−41^m^2^/V^2^. This translates to a *γ* value of 3.9 ± 0.5 × 10^−33^esu, which is just within error of the *γ* value reported by Tuer et al (2010) (1.8 ± 1.4 × 10^−32^esu) [21]. The *γ* value of eosin Y is close to that of *β*-carotene (4.9 0.8 10^−41^m^2^ /V^2^), which is a known THG harmonophore [40, 42, 44].

The H-stained tissue shows similar MPF intensity from ECM compared to the unstained tissue (Fig. 1d). The SHG intensity of the H-stained tissue is similar to that of the unstained tissue (Fig. 1e), while THG appears to be strong, highlighting the cell nuclei (Fig. 1f). The intense THG originates from the hemalum aggregates formed in the nuclei during H staining [21]. In addition, a weak THG signal is observed in H-stained ECM. The signal is more than one order of magnitude weaker than in the nuclei, but it is about the same intensity as that from the E-stained collagen fibers (see inset of Fig. 1f). In contrast, the THG signal of H-stained collagen fibers appears stronger than in unstained tissue (compare inset in Fig. 1f with panel l) indicating that some hemalums remain in the tissue stroma after washing step with water [45].

The H&E-stained tissue shows slightly lower MPF intensity compared to the E-stained collagen (Fig. 1a) which is due to eosin fluorescence quenching by hemalums [21]. The SHG intensity is about two times lower in H&E-stained tissue (Fig. 1b) than in the unstained tissue, suggesting a significant effect from H&E staining on the SHG signal. A strong THG signal is observed from the cell nuclei in H&E-stained tissue, similar to the nuclei in H-stained cells (Fig. 1c). The THG signal from the H&E-stained collagen fibers appears lower than from the cell nuclei, however, it is more than one order of magnitude stronger compared to the THG signal from the stroma in H-stained or E-stained collagen fibers (compare inset in Fig. 1c with insets in panel f and panel i). Therefore, THG microscopy provides a convenient method for visualizing not only the nuclei but also ECM in H&E-stained tissues. Note that the *γ* values of hemalums and hemalum and eosin complexes could not be determined due to aggregation at neutral pH [21].

In summary, multimodal imaging reveals that the strongest MPF originates mostly from eosin and is partially quenched by hemalums in H&E-stained tissue. The SHG originates from collagen, and H&E staining significantly decreases the SHG signal intensity, while H staining and E staining have negligible influence on the SHG. The THG originates from hemalum aggregates in the cell nuclei and it is enhanced when both hemalums and eosin stains interact [21]. The hemalums and eosin interact within the ECM and exhibit an order of magnitude stronger THG signal in the stroma of H&E-stained tissue compared to the stroma of H-stained or E-stained tissue. The THG is relatively weak in unstained tissue and does not contribute significantly to the signals of stained tissues.

### 3.2. Photobleaching of nonlinear signals in differently stained tissues

The influence of hematoxylin and eosin staining on the nonlinear signals is investigated by reducing the stain concentration in the tissues via photobleaching. To induce photobleaching, small areas of the samples are scanned with actinic light for 90 seconds having an increased laser power of 30 mW at the sample. Subsequently, MPF, SHG and THG images are recorded from larger areas with lower laser power to visualize the bleached and unbleached regions (Fig. 2).

**Fig. 2.**
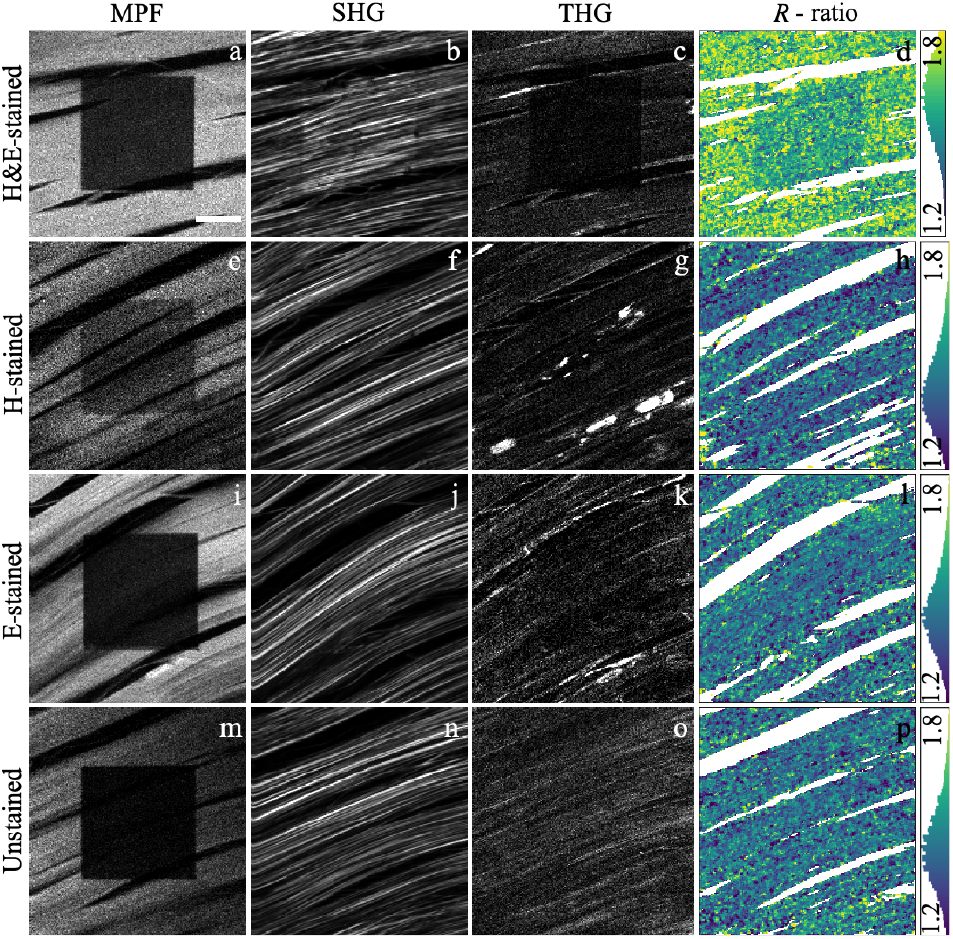
Bleaching of nonlinear signals in H&E-stained (a-d), H-stained (e-h), E-stained (i-l) and unstained (m-p) tissues. MPF (a, e, i, m), SHG (b, f, j, n), THG (c, g, k, o) images and SHG *R* ratio maps (d, h, l, p) are shown in corresponding columns. The bleached area is visible in the center of the images. The same area is imaged with MPF, SHG and THG for each of the corresponding tissue staining types. Scale bar 20 µm.

After photobleaching with the chosen actinic laser powers and scanning durations, between 10 and 47 % of the initial fluorescence intensity remains (see Table 1). Photobleaching is a complex process leading to a concentration reduction of fluorophores, or/and generation of quenchers that reduce fluorescence yield. In this analysis, we assume that photobleaching primarily decreases the concentration of fluorophores. The photobleaching of unstained tissue reduces the MPF to 10 % of the unbleached intensity level. Autofluorescent molecules that can emit within the applied filter transmission range include collagen, elastin, NADH and flavins [46]. The MPF of E-stained tissue is bleached to 16 % of the initial level. The fluorescence of H-stained tissue is bleached to 47 %, while H&E-stained tissue bleached to 22 % of the initial level. The change in the bleaching levels approximately corresponds to the change in concentration of the fluorophores in respective tissues. The bleaching of autofluorescence contributes to the overall bleaching of the stained tissues, particularly affecting the H-stained sample due to low fluorescence yield of hemalums.

**Table 1.** Fractions of average MPF, SHG and THG intensity remaining after bleaching for unstained, E-stained, H-stained and H&E-stained tissue shown in Fig. 2, and SHG *R* ratio values of unbleached and bleached areas.

| | MPF | SHG | THG | $R$ (unbleached) | $R$ (bleached) |
| --- | --- | --- | --- | --- | --- |
| H&E-stained | 0.22 | 1.51 | 0.30 | 1.64 | 1.56 |
| H-stained | 0.47 | 1.00 | 0.70 | 1.42 | 1.40 |
| E-stained | 0.16 | 1.00 | 0.57 | 1.44 | 1.42 |
| Unstained | 0.10 | 1.00 | 1.00 | 1.43 | 1.43 |

The SHG images of bleached and unbleached areas show that the SHG intensity does not change in unstained, H-stained and E-stained tissues (Fig. 2n, j, f). Interestingly, SHG increases (i.e., antibleaches) in the areas treated with actinic light in H&E-stained tissue (Fig. 2b). The antibleaching of SHG was previously observed in widefield SHG microscopy of tendon tissue [27]. Its origin is not fully understood and was previously attributed to the reabsorption of SHG signal by the stains. By reduction of the stain concentration, the SHG reabsorption decreases.

The THG signal intensity is not affected by the bleaching in unstained tissue. In contrast, the THG signal is bleached to 57 % in E-stained tissue, and to 70 % in H-stained tissue, compared to the unbleached area (Table 1). The largest bleaching effect reaching to 30 % of the unbleached THG level is observed in H&E-stained tissue. The average intensity ratios of bleached and unbleached areas (Table 1) show that stain molecules contribute to THG, and the biggest contribution appears in the H&E-stained tissue. The THG images also reveal that hemalum and eosin molecules interact in H&E-stained tissue, resulting in higher THG signal in the nuclear regions and, particularly, in the ECM collagen areas, compared to H-stained, E-stained and unstained tissues (Fig. 1).

### 3.3. Organization of harmonophores in tendon tissue revealed by SHG polarimetry

P-SHG imaging is employed to investigate the ultrastructural organization of the stained collagen in the bleached and unbleached areas. The *R* ratio maps are shown in Fig. 2 for each staining case, and the average *R* ratio values of unbleached and bleached areas are listed in Table 1. The *R* ratio values appear to be the same for unstained, H-stained and E-stained tissues. The unbleached area of H&E-stained tissue has a higher *R* ratio compared to the unstained, H-stained and E-stained tissue (see Fig. 2). Bleached H&E-stained tissue has a lower *R* ratio but it is still higher that in the other samples (Table 1). The SHG signal intensity exhibits no bleaching in the H-stained, E-stained and unstained samples, and undergoes antibleaching in the H&E-stained sample. This antibleaching effect of SHG in the H&E sample, the increased *R* ratio due to staining and the *R* decrease after bleaching suggest that hemalums and eosin together affect the second-order susceptibility tensor of the tendon tissue. Such an effect can be caused by a noncentrosymmetric (polar) arrangement of hemalum and eosin molecules in the collagenous tissue. The hemalums and eosin may form complexes that arrange with *C*_6_ symmetry along the collagen fibers. The polar organization of the dyes results in the increase of *R* ratio.

### 3.4. Organization of harmonophores in tendon tissue revealed by THG polarimetry

The ultrastructural organization of hematoxylin and eosin molecules in histology tissue is further studied using polarization-resolved THG microscopy. A *C*_6*v*_ THG PIPO model is fitted to the datasets to obtain the orientation of the cylindrical axis and two susceptibility tensor component ratios 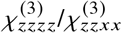 and 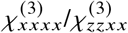. An isotropic distribution of harmonophores corresponds to 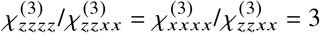 ratios [33, 47], and if molecules are partially aligned the ratios deviate from the isotropic values [31]. Fit values for all samples are presented in Fig. 3. The figure shows THG intensity images (a, e, i, m), the maps of susceptibility ratios 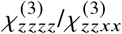 and 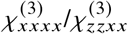 (b, f, j, n), and the *δ* orientation maps of the cylindrical symmetry axis with respect to Z-axis of the laboratory frame (d, h, l, p). PIPO plots show THG intensity as a function of the incident and outgoing polarization orientation and are presented in Fig. 4 for the regions of interest (ROIs) indicated by rectangles in Fig. 3 a, e, i, and m.

**Fig. 3.**
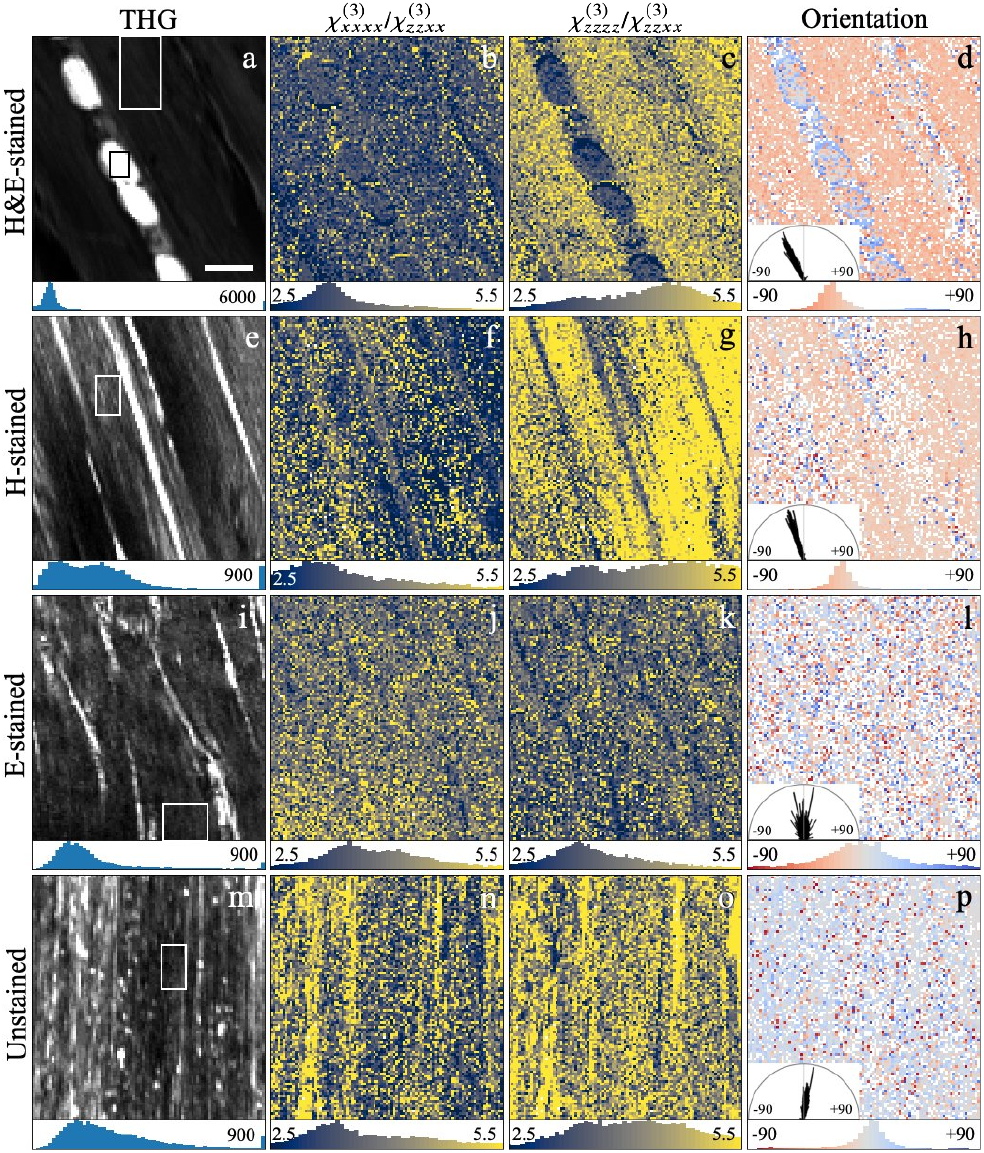
THG intensities and PIPO fit results of H&E-stained (a-d), H-stained (e-h), E-stained (i-l) and unstained RTT tissue (m-p). THG intensity images (a, e, i, m), maps of 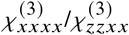 (b, f, j, n) 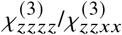 (c, g, k, o) ratios and orientation *δ* (d, h, l, p) are shown in the columns. Pixel value histograms and corresponding color maps are shown below the images. Polar histograms are shown in panels d, h, l, p. Only one half of the polar space is needed to show the unsigned orientation of intensity. ROIs for 2D PIPO plots (shown in Fig. 4) are marked by rectangles (a, e, i, m). The maximum count value is set so that <5% of pixels can exceed it in panels a, e, i, and m. Scale bar 10 µm.

**Fig. 4.**
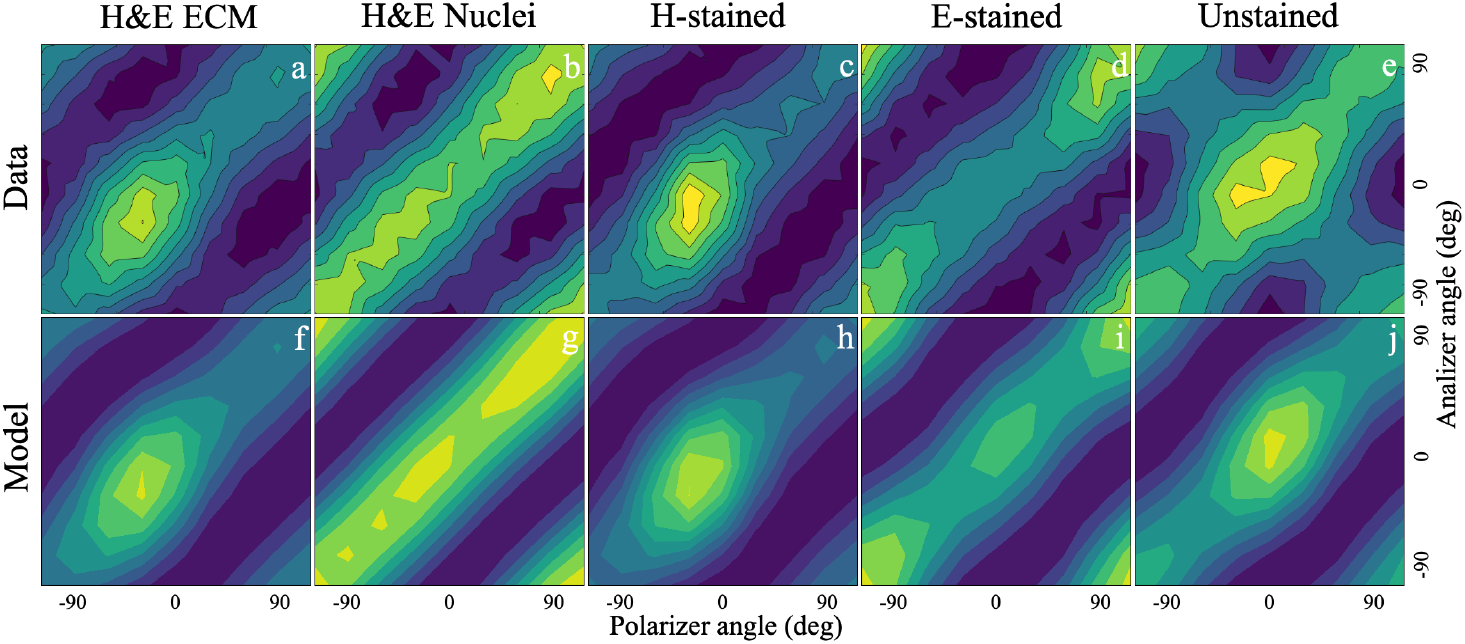
2D THG PIPO data (a-e) and the best-fit *C*_6_ model (f-j) plots. The data taken from ROIs shown in Fig. 3 a, e, i, m and the fits are for H&E-stained ECM (a, f) and nuclei (b, g), H-stained (c, h), E-stained (d, i) and unstained (e, j) tissues, respectively.

The unstained tissue gives low THG signal, which can be seen in Fig. 1o. Nevertheless, it is possible to collect enough signal for PIPO analysis, albeit with noisy ratio maps (see Fig. 3m-p). The delta map shows a dominant cylindrical axis orientation along the collagen fibers with narrow distribution seen in the polar histogram (Fig. 3p). The values of 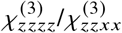 are mostly distributed between 4 and 5, while 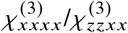 values peak around 3.8. The PIPO plots of selected ROIs show well defined distributions of THG intensity along the collagen fibers (Fig. 4e, j). The PIPO plots of the unstained sample can be used as a reference to compare against the modifications of 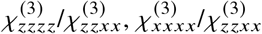 and cylindrical axis orientation *δ* as a result of staining with hemalums and eosin.

The THG intensity of E-stained tissue is comparable to unstained tissue. The peak values of 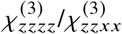 and 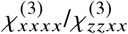 distributions are slightly larger than 3 (Fig. 3j, k) indicating that the eosin dye has a broad orientation distribution approaching isotropic organization, although the molecular organization still displays a cylindrical symmetry. 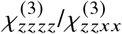 has lower values than 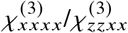, suggesting that eosin molecules are arranged cylindrically, with their nonlinear dipoles oriented at large angles relative to the cylindrical symmetry axis. The cylindrical axis has preferential orientation along the collagen fibers with a broader *δ* value distribution than for the unstained sample (compare Fig. 3l and p). The PIPO plot of the selected ROI with tendon fibers shows slight deviation from isotropic arrangement (Fig. 4d, i). The THG maximum intensity position along the diagonal of PIPO plot has close to 90^°^ shift with respect to unstained fibers (compare Fig. 4 d, and e), indicating that eosin nonlinear dipoles have a preferential arrangement at large angles with respect to the collagen fiber axis.

The H-stained RTT tissue shows large THG PIPO differences (Fig. 3e-h) compared to the E-stained tissue (Fig. 3i-l). The 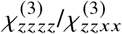 of the fiber regions has values above 5 and the 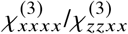 values are around 3 (Fig. 3f, g), showing that nonlinear dipoles of hemalums orient with small angles to the cylindrical axis. The orientation of the cylindrical axis of ordered hemalums is along the collagen fibers (Fig. 3h). The PIPO plot (Fig. 4c, h) of the ROI from the stroma region (indicated with rectangle in Fig. 3e) shows high 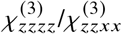 values and the peak of THG intensity occurs with polarization orientations along the collagen fibers. It is interesting to note that although hemalums accumulate mostly in the nuclei, some molecules remain in the ECM after washing step during the preparation procedure [45], and align at small angles with respect to collagen fibers. The edges of collagen fibers have lower 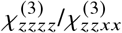 values approaching 3, which represents lower order of hemalums at those locations.

Two types of PIPO response can be found in H&E-stained samples. The nuclei regions have predominantly isotropic THG response characterized by 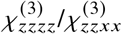 and 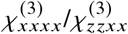 values around 3 (Fig. 3b, c). The PIPO plot for the nuclear ROI shown with black frame in Fig. 3a is given in Fig. 4b, g. It can be seen that for any incident laser polarization angle the maximum THG emission intensity is achieved correspondingly at the same output angle. The origin of isotropic THG signal is due to randomly organized hemalum and eosin aggregates in the nuclei [21]. On the other hand, the H&E-stained sample shows the anisotropic THG polarization response originating from the stroma regions with 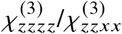 values well above 4 and 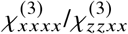 values around 3 (Fig. 3b, c). The higher values of 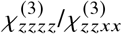 compared with 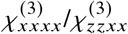 indicate that the molecular dipoles of the dyes are oriented close to the cylindrical symmetry axis [31]. The cylindrical axis aligns preferentially along the collagen fibers. The PIPO plots (Fig. 4a, f) are also clearly different in the fiber ROI (shown with white frame in Fig. 3a) compared to that of the nuclei (Fig. 4b, g). The THG signal in the fiber ROI is strongest when the incident light is polarized at 30^°^ from the laboratory Z axis, i.e., along the collagen fibers (Fig. 4a, f).

P-THG shows that organization of nonlinear dipoles can be approximated by a cylindrical symmetry with axis orientation along the collagen fibers for all four sample types. The nonlinear dipoles are arranged in a cone with a half-angle less than 45^°^ for unstained, H-stained and H&E-stained samples. The eosin stain has a broad orientation distribution of nonlinear dipoles with the cone half-angle higher than 45^°^. The difference in 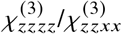 and 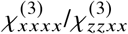 ratios for the H&E sample, compared to H-stained and E-stained tissues, provides evidence for an interaction of hemalums and eosin. The hemalums and eosin interact and align along the collagen fibers with a cone half-angle larger than in H-stained, and smaller than in E-stained sample.

### 3.5. Orientation of hematoxylin and eosin in tendon tissue revealed by FDMLD

The orientations of hemalums and eosin in tendon tissue are also studied with polarimetric MPF microscopy. Fig. 5 shows FDMLD of vertically oriented tendon in unstained (Fig. 5h), E-stained (Fig. 5g), H-stained (Fig. 5f) and H&E-stained (Fig. 5e) tissues. The unstained tissue shows isotropic fluorescence with FDMLD values distributed around 0. The E-stained tissue has higher fluorescence intensity when excited with vertically polarized light, and shows FDMLD distribution peaking at 0.21. The results indicate that nonlinear absorption dipoles of eosin are preferentially oriented along the collagen fibers. The vertically aligned H-stained tendon exhibits higher FDMLD values peaking at 0.32. It provides evidence that nonlinear absorption dipoles of hemalums are aligned closer to the fiber axis of collagen compared to the nonlinear absorption dipoles of eosin. In H&E-stained tissue, FDMLD values (peaking at 0.21) are distributed very similar to E-stained tissue. The eosin fluorescence is quenched by hemalums in hemalum-eosin complexes [21]. Therefore, FDMLD preferentially detects the fraction of unquenched eosin. The orientation distribution of unquenched eosin is very similar in H&E- and E-stained tissues.

**Fig. 5.**
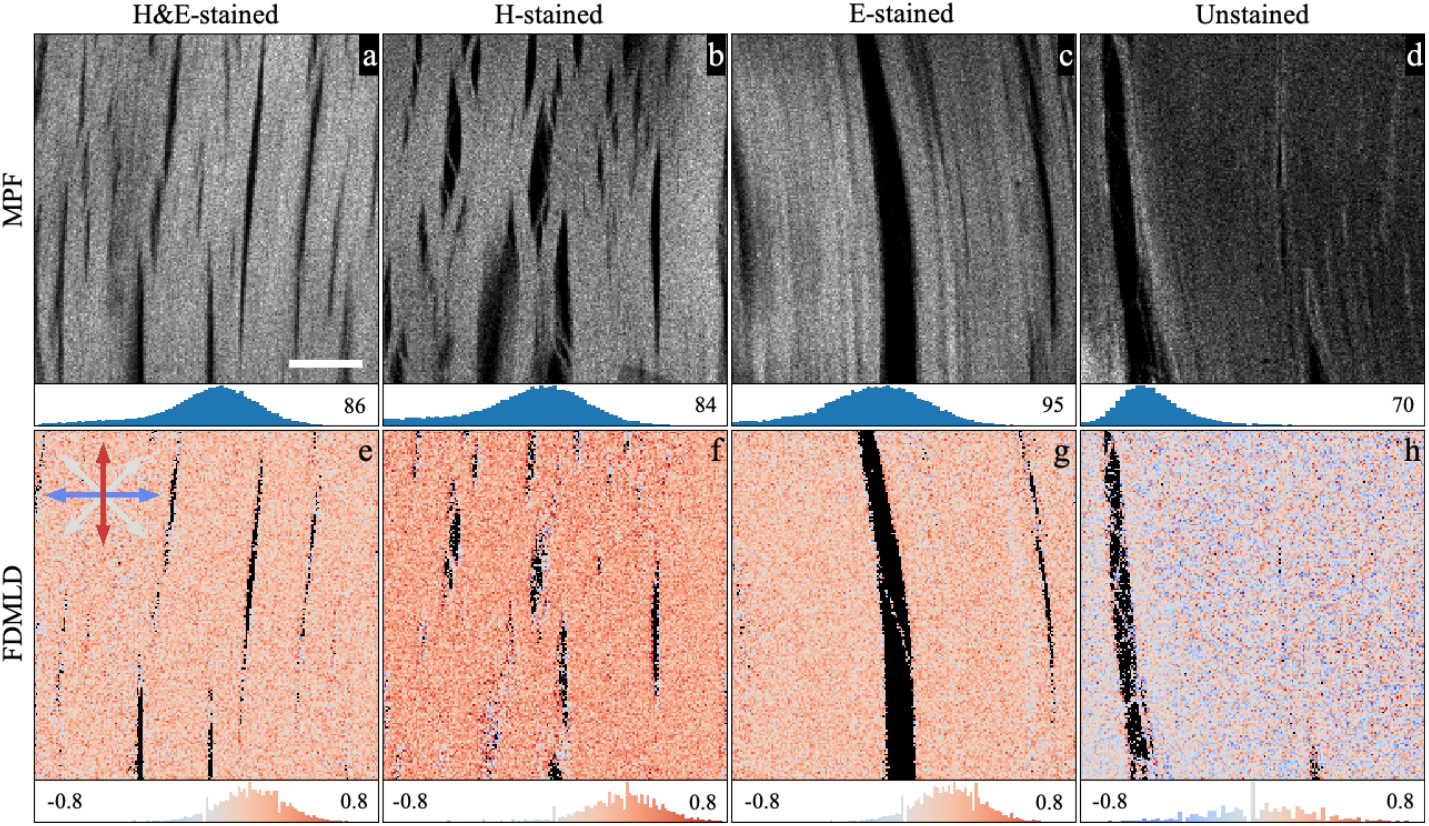
MPF intensity images (a-d) and FDMLD maps (e-h) of H&E-stained (a, e), H-stained (b, f), E-stained (c, g) and unstained (d, h) tissues. Similar fluorescence intensities are obtained by using different laser powers as follows: for H&E stained tissue - 1.5 mW (a), H-stained - 7 mW (b), E-stained - 1 mW (c) and unstained - 8 mW (d). The color code of orientations of nonlinear absorption dipoles is shown with arrows in (e). The MPF intensity and FDMLD value distributions with color code are shown below the corresponding images. Scale bar 20 µm.

## 4. Discussion

### 4.1. Influence of hemalums and eosin on nonlinear optical properties of collagen tissue

Polarimetric SHG, THG and MPF microscopy reveals that hemalum and eosin stains align with the collagen fibers. This is in line with the increased birefringence of H-stained, E-stained and H&E-stained collagen fibers [45]. Based on the SHG and THG PIPO models the molecular organization of the aligned stains can be approximated by cylindrical symmetry with the axis parallel to the collagen fiber axis. Eosin Y is loosely aligned at larger angles with respect to the cylindrical axis, while hematoxylin has a narrower orientation distribution with nonlinear dipoles aligned closer to the cylindrical fiber axis. Both dyes, when used separately, do not significantly influence the second-order susceptibility tensor component ratio *R*, indicating that the dye molecules orient with *C*_6*h*_ symmetry configuration, i.e. with a flip-flop arrangement along the collagen fibers.

When both hemalum and eosin stains are used together, the dyes interact leading to a decrease in SHG intensity of collagen tissue, increase in THG intensity of the ECM, and eosin fluorescence quenching by hemalums [21]. The polarimetric THG of H&E-stained tissue shows the alignment of hemalums and eosin molecules along the collagen fibers, which differs from the alignment of hemalums or eosin when stained separately. This is in line with the presence of birefringence in H&E-stained fibers, which appears somewhat less pronounced than with H-staining or E-staining alone [45]. Most interestingly, the higher *R* ratio in H&E-stained tissue and the ratio decrease upon bleaching shows a noncentrosymmetric arrangement of the hemalums and eosin with *C*_6_ or *C*_6*v*_ symmetry that follows the polarity of collagen. The fact that SHG intensity is lower in H&Estained tissue and increases upon bleaching indicates that the nonlinear susceptibility components of the ordered dyes have opposite phase shift compared to the nonlinear susceptibility of collagen. The phase differences may appear due to the spectral differences of nonlinear absorption of collagen, hemalums and eosin, and due to the differences in geometrical arrangement of nonlinear dipoles for collagen and the dyes. The increase in SHG intensity upon bleaching has been observed with wide-field SHG microscopy of H&E-stained tissues, and was interpreted as being due to reabsorption of collagen SHG by the H&E stains [27]. This might still partially be the case, however, the higher susceptibility ratio *R* in the H&E-stained tissues and its reduction upon photobleaching points to the second order susceptibility contribution due to the presence of noncentrosymmetrically aligned stain molecules with *C*_6_ or *C*_6*v*_ symmetry in the collagenous tissue. The reported changes in SHG circular dichroism upon H&E staining [28] point to the chiral *C*_6_ organization of hemalums and eosin along the collagen fibers in the ECM.

### 4.2. Molecular organization of hemalums and eosin in the extracellular matrix

The H&E staining procedure involves first staining with hemalums at low pH, followed by the blueing step which raises the pH due to washing the specimens with tap water. Binding of hematein to the collagen fibers either directly or via coordinating with aluminum cations leads to the alignment of hemalums along the collagen fibers as shown with polarimetric THG and MPF microscopy of H-stained tissue (Figs 3, 4, 5). The H-staining also increases the birefringence of collagen fibers [45]. Collagen is comprised of a repeated sequence of glycine, proline and hidroxyproline and contains negatively charged aspartic and glutamic acids as well as positively charged lysine, arginine and histidine amino acids [48]. The aspartic and glutamic acids, being negatively charged at neutral pH, can interact with positive hemalum ions. Additionally, hydroxyproline is a polar amino acid and can efficiently interact with hemalums via hydrogen bonding providing the observed alignment along the collagen fibers. The hemalums are present in various forms at low pH e.g. Hm, HmAl^2+^, HmAl^+^, and minor amount of HmAl^0^, and interact with collagen fibers during staining [21]. By raising pH to neutral during the blueing step, HmAl^0^ becomes the dominant form and accumulates in the nuclei [21], while the fluorescent HmAl^2+^ form aligns along the collagen fibers in the ECM as shown by MPF microscopy. Probably, HmAl^1+^ and HmAl^0^ also align along the collagen fibers and together with HmAl^2+^ contribute to the polarimetric response of THG. On the other hand, the organization of hemalums along the collagen fibers does not change the *R* ratio and SHG intensity of collagen, as revealed by P-SHG of H-stained tissue. Thus, it is likely that HmAl^2+^, HmAl^1+^ and HmAl^0^ each having two isomeric forms [49] do not influence the SHG response due to their *C*_6*h*_ organizational symmetry i.e. having a flip-flop alignment of harmonophores along the collagen fibers.

When anionic eosin Y, having up to two negative charges at neutral pH [50, 51], is applied to the unstained tissue, it interacts with positively charged lysine, arginine and potentially histidine amino acids of collagen [52]. Eosin Y can also interact with collagen via Van der Waals forces [53]. Birefringence is enhanced in E-stained collagen fibers [45] demonstrating dye ordering. The eosin dye partially aligns along the collagen fibers as demonstrated by FDMLD (Fig. 5). P-THG also shows a partial alignment of eosin with cylindrical axis along the collagen fibers and nonlinear dipoles arranged at large angles with respect to the fiber axis (Fig. 3). This demonstrates a loose alignment of eosin molecules around the collagen fibers. In addition, P-SHG of collagen in E-stained tissue appears to be the same as in the unstained tissue. Therefore, the polarimetric results show that eosin Y molecules arrange with cylindrical *C*_6*h*_ (flip-flop) symmetry along the collagen fibers.

When eosin staining is applied to the tissue stained with hemalums, the eosin anions interact electrostatically with the hemalum cations and the positive amino acids of collagen. The fluorescence intensity decrease due to quenching [21] and the THG increase in the ECM of H&E-stained tissue provide evidence for complex formation between the dyes (Fig. 1). The hemalum-eosin complexes do not produce significant new absorption bands in the H&E-stained tissue compared to the spectra of H-stained and E-stained tissue [21]. However, the fluorescence quenching of eosin by hemalums shows that hemalum and eosin molecules are positioned within the Förster radius, probably via coordination of the aluminum ions. Interestingly, the SHG intensity decrease upon H&E staining, the antibleaching effect and the alteration of the *R* ratio reveal a likely *C*_6_ organization of hemalums and eosin, which form complexes that show alignment polarity preferentially in one direction along the cylindrical axis of the collagen fibers. The influence of H&E staining on the SHG-CD [28] also points to the chiral organization of hemalum-eosin complexes along collagen fibers. The noncentrosymmetric organization of the complexes follows the polarity of collagen. Collagen is comprised of the repeated triplet units containing glycine, proline and hydroxyproline amino acids [54]. The hemalum and eosin complexes likely form hydrogen bonds with the hydroxyl groups of hydroxyprolines, and also interact with the charged amino acid residues leading to the polar alignment of the dyes. The Van der Waals forces may also play a role in the noncentrosymmetric alignment of hemalum and eosin complexes. The elucidation of the main factors leading to the noncentrosymmetric alignment of hemalum and eosin complexes along collagen fibers requires further investigation. However, the polar alignment of the hemalum and eosin complexes has to be kept in mind when analyzing the polarimetric linear and nonlinear microscopy data in H&E tissue histopathology.

## 5. Conclusions

Polarimetric THG and MPF microscopy reveals that the nonlinear dipoles of hemalums in H-stained tissue are oriented at small cone angles to the cylindrical symmetry axis of the fibers, while the nonlinear dipoles of eosin are distributed more broadly from the fiber axis in E-stained tissue. In H&E-stained tissue, the hemalum and eosin molecules have an intermediate spread from the fiber axis, falling between the narrow distribution of hemalums in H-stained tissue and the broader distribution of eosin in E-stained tissue.

The H&E staining of histology tissue leads to the noncentrosymmetric organization of hemalum and eosin complexes with *C*_6_ symmetry, having polar alignment along the collagen fibers. The polar arrangement of the dyes decreases the SHG intensity and increases the achiral nonlinear susceptibility ratio *R* compared to the unstained collagen. The treatment with intense actinic light leads to the SHG intensity increase (antibleaching) and *R* ratio decrease in the ECM. These effects have to be considered in SHG microscopy of H&E-stained tissue histology. In contrast, the organization of hemalums in H-stained tissue and eosin in E-stained tissue both follow *C*_6*h*_ symmetry with cylindrical axis aligned along the collagen fibers. Therefore, the staining does not affect the SHG intensity and the *R* ratio in H-stained and E-stained histology tissues.

These polarization effects have to be considered more comprehensively in the multiphoton microscopy of H&E-stained tissue when laser polarization is not accounted for in the imaging process. “Intensity-only” imaging is typically performed with a fixed incident laser polarization and the detected intensity depends on the orientation of collagen fibers with respect to this polarization. First, H&E samples exhibit a higher SHG *R* ratio, which decreases with the exposure, while the SHG intensity itself increases with the exposure. This results in changes of SHG image morphology over time during prolonged exposure. Second, MPF of eosin is polarization-dependent, leading to varying intensities depending on the laser polarization and the orientation of the sample. Third, autofluorescence of unstained tissue is not sensitive to the laser polarization, but SHG and THG signals are dependent on the laser polarization and the orientation of the fibers. Therefore, polarization effects have to be taken in consideration when working with hematoxylin and eosin stained tissues.

## Acknowledgments

We thank Dr. Rotomskis, Vilnius University, for fruitful discussions, and “Light Conversion” for providing the laser. This work is supported by the Natural Sciences and Engineering Research Council of Canada (NSERC) (RGPIN-2024-06356 for VB, and RGPIN-2018-05444 for DT), Research Council of Lithuania (LMT) (P-MIP-24-644 for VB), the European Regional Development Fund with LMT (01.2.2.-LMT-K-718-02-0016 for VB), and Canada Foundation for Innovation (John R. Evans Leaders Fund 37749 for DT), Research Nova Scotia 1868 for DT, and Saint Mary’s University for DT.

## Data availability

Data is available from V.B. upon reasonable request.

